# Sub-atomic resolution crystal structures reveal conserved geometric outliers at functional sites

**DOI:** 10.1101/713818

**Authors:** Saara Laulumaa, Petri Kursula

## Abstract

P2 is a peripheral membrane protein of the vertebrate nervous system myelin sheath, having possible roles in both lipid transport and 3D molecular organization of the multilayered myelin membrane. We extended our earlier crystallographic studies on human P2 and refined its crystal structure at an ultrahigh resolution of 0.72 Å in perdeuterated form and 0.86 Å in hydrogenated form. Characteristic differences in C-H…O hydrogen bond patterns were observed between extended β strands, kinked or ending strands, and helices. Often, side-chain C-H groups engage in hydrogen bonding with backbone carbonyl moieties. The data highlight several amino acid residues with unconventional conformations, including both bent aromatic rings and twisted guanidinium groups on arginine side chains, as well as non-planar peptide bonds. In two locations, such non-ideal conformations cluster, providing proof of local functional strain. Other ultrahigh-resolution protein structures similarly contain chemical groups breaking planarity rules. For example, in SH3 domains, a conserved bent aromatic residue is observed near the ligand binding site. FABP3, belonging to the same family as P2, has several side chains and peptide bonds bent exactly as those in P2. We provide a high-resolution snapshot on non-ideal conformations of amino acid residues under local strain, possibly relevant to biological function. Geometric outliers observed in ultrahigh-resolution protein structures are real and likely relevant for ligand binding and conformational changes. Furthermore, deuteration of protein and/or solvent are promising variables in protein crystal optimization.

## 1. Introduction

As biological molecules work in mainly aqueous environments, details of protonation and hydrogen bonding are crucial to understanding protein function at the atomic level. Perdeuterated proteins can be used in neutron crystallography for accurate determination of hydrogen positions and protonation states. On the other hand, it is possible that perdeuteration affects either the structure or the ligand-binding properties of the protein – or both [1]. Using ultrahigh-resolution X-ray crystallography, many hydrogen atom positions can also be determined, but relatively low levels of local disorder will already make H atoms invisible in electron density. Ultrahigh resolution also allows to detect structural anomalies, both in the peptide backbone and in the side chains, due to the possibility of X-ray data outweighing geometrical restraints during refinement.

The crystal structures of only 10 different proteins have been refined at a resolution of at least 0.75 Å, and 8 of these proteins are larger than 100 amino acid residues. Three human proteins have been refined at 0.75-Å or higher resolution: cyclophilin G [2], aldose reductase [3], and the PDZ domain of syntenin [4]. Highest resolution for any protein structure has been achieved for crambin, at 0.48 Å [5], and the perdeuterated form of *Pyrococcus furiosus* rubredoxin has been refined at 0.59-Å resolution [6].

P2 is a myelin-specific protein of the nervous system, which acts as a peripheral membrane protein within the myelin sheath. P2 belongs to the FABP family, and it is likely to function in lipid transport during myelination. P2 has a role in maintaining lipid homeostasis in the developing nervous system [7]. Mutations in the gene encoding P2 are linked to Charcot-Marie-Tooth disease [8–12]. We have previously determined the atomic-resolution 0.93-Å structure of human P2 [13], providing accurate insights into its ligand-binding determinants. The positions of many hydrogen atoms were also defined within the internal hydrogen-bonding network, and one of the fatty acid-coordinating arginine residues was shown to be in the neutral deprotonated form.

We set out to take advantage of modern synchrotron radiation beamlines and the exceptional diffraction properties of human myelin P2 protein crystals to get an even deeper insight into the structural details of this peripheral membrane protein. The human P2 structure was refined at 0.72-Å resolution, using data collected from perdeuterated P2 (d-P2), and a number of intriguing details concerning protein structure are revealed. In addition, the structure of hydrogenated P2 (h-P2) was refined to 0.86-Å resolution using new data. The complementary use of these data gave directions to analyzing conformational abnormalities in other proteins and protein families, highlighting conservation of conformational strain in locations possibly important for ligand binding, catalysis, and/or conformational changes.

## 2. Results and Discussion

### 2.1. Overall structure and quality

Based on CD spectra, the folding of the perdeuterated human P2 was practically identical to the hydrogenated protein (Figure 1a). Thermal unfolding assays indicated d-P2 to be stable in aqueous solution, with a melting temperature of +60 °C, which is similar to that observed for h-P2 earlier [13,14]. Upon crystallization experiments for neutron diffraction [15], large d-P2 crystals were also tested on synchrotron X-ray beamlines and observed to give sub-atomic resolution diffraction patterns [16,17].

**Figure 1.**
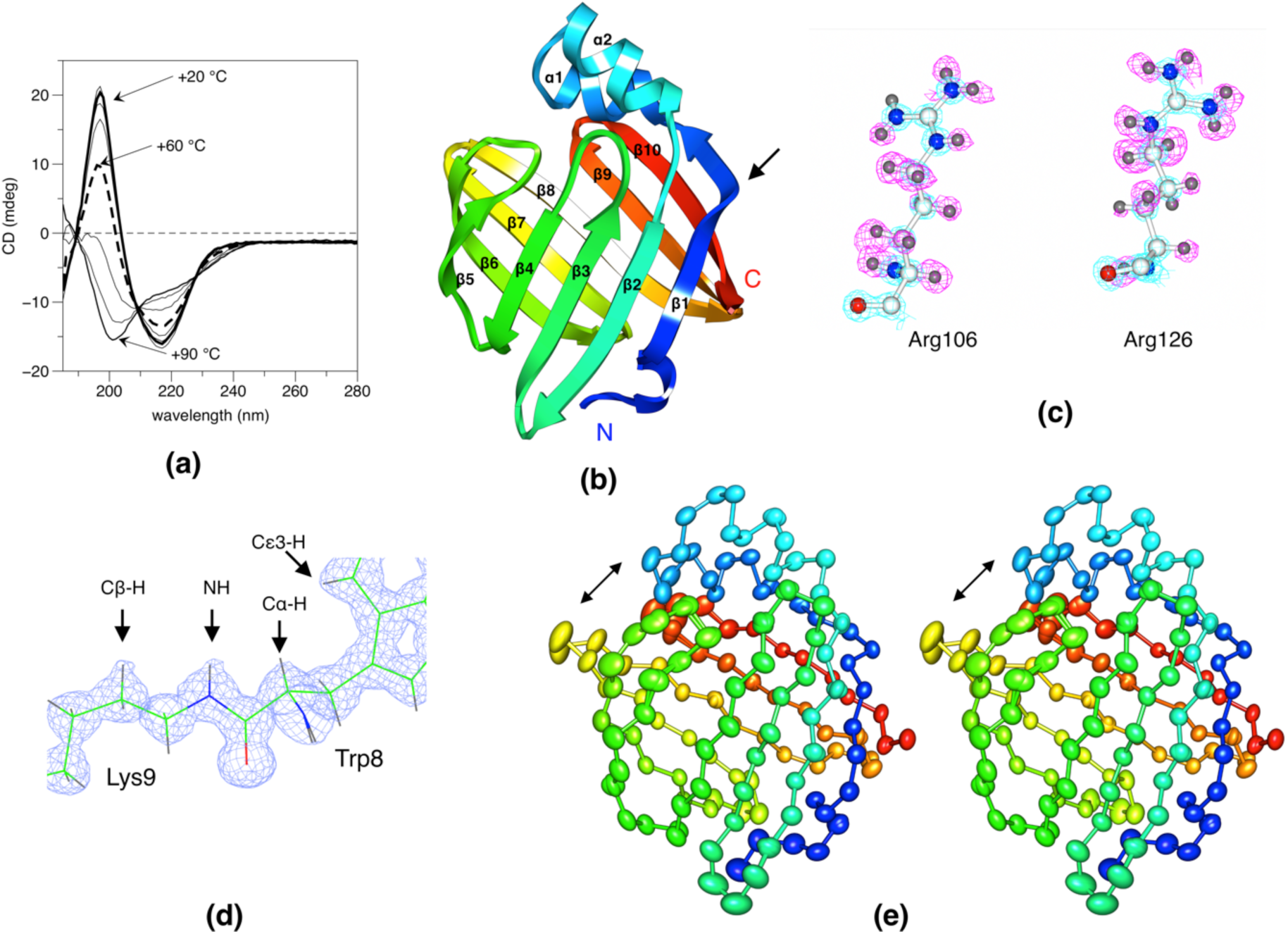
Structural properties of deuterated P2. (**a**) CD spectra as a function of temperature from +20 to +90 °C. For clarity, only spectra at 10 °C intervals are shown. (**b**) Overall structure of human P2. The secondary structures and the termini are labelled. The arrow points at the bump in strand β1. (**c**) Quality of electron density for the two Arg residues, which coordinate the bound fatty acid. The 2F_o_-F_c_ map (blue) is contoured at 4.0 σ and the F_o_-F_c_ map (magenta) at 2.0 σ. Note how Arg106 is deprotonated, *i*.*e*. in a neutral form, as we also reported prevously [13]. The maps are for h-P2 at 0.86-Å resolution, calculated without hydrogen atoms. (**d**) The 2F_o_-F_c_ map for d-P2 shows clear bumps at the positions of well-defined deuterium atoms, indicating very high quality of the diffraction data. (**e**) Analysis of the anisotropic displacement parameters suggests open-close motions (arrows) at the portal region even in the crystal state (stereo view).

The structure of perdeuterated human P2 was refined at 0.72-Å resolution with good statistics (Table 1). The overall structure is very similar to our earlier structure of the hydrogenated protein (Figure 1b), but the space group is different. The packing in the new space group, in fact, has caused some disorder despite the high resolution of X-ray diffraction (data not shown).

**Table 1.** Data processing and structure refinement. The values in parentheses refer to the highest-resolution shell.

| Sample | d-P2 | h-P2 |
| --- | --- | --- |
| Space group | C2 | P41212 |
| Unit cell dimensions | a=112.18 Å, b=36.21 Å, c=31.11 Å,<br>$\alpha=\gamma=90^\circ$, $\beta=97.03^\circ$ | a=b=57.93 Å, c=101.32 Å,<br>$\alpha=\beta=\gamma=90^\circ$ |
| Wavelength (Å) | 0.7443 | 0.8266 |
| Resolution range (Å) | 50-0.72 (0.74-0.72) | 30-0.86 (0.88-0.86) |
| $\langle I/\sigma(I) \rangle$ | 15.8 (0.9) | 14.7 (1.1) |
| R<sub>sym</sub> (%) | 3.6 (121.4) | 6.3 (169.9) |
| R<sub>meas</sub> (%) | 3.9 (143.5) | 6.8 (183.0) |
| Completeness (%) | 91.0 (53.2) | 94.4 (87.1) |
| Redundancy | 4.7 (3.3) | 7.2 (7.0) |
| CC<sub>1/2</sub> (%) | 99.9 (40.5) | 99.8 (43.3) |
| Wilson B factor (Å<sup>2</sup>) | 9.0 | 11.3 |
| R<sub>cryst</sub> (%) | 10.4 | 9.9 |
| R<sub>free</sub> (%) | 11.1 | 11.8 |
| rmsd bond lengths (Å) | 0.020 | 0.020 |
| rmsd bond angles (°) | 1.8 | 1.9 |
| Average B factor (Å<sup>2</sup>); protein,<br>ligand, solvent | 10.7, 14.1, 21.4 | 11.0, 11.6, 21.6 |
| Ramachandran favoured/allowed<br>(%); Molprobity score (percentile) | 100/100; 1.30 (85<sup>th</sup>) | 99.2/100; 1.25 (86<sup>th</sup>) |
| Mean anisotropy; protein, ligand, solvent | 0.41 ± 0.13, 0.41 ± 0.14, 0.40 ± 0.15 | 0.49 ± 0.15, 0.46 ± 0.12, 0.41 ± 0.16 |
| PDB entry | 6S2M | 6S2S |

In the ligand-binding cavity, the new crystal form has more disorder than observed in the previous 0.93-Å structure, and hence, less hydrogen atoms are observable *e*.*g*. within the water-mediated hydrogen-bonding network. It is possible that deuteration weakens the interactions between the protein and the ligand/water molecules in the cavity such, that they become more dynamic. On the other hand, the protein structure is of very high quality and fascinating details become visible at this sub-atomic resolution.

In addition to d-P2, an ultrahigh-resolution structure of h-P2 was also refined, at 0.86-Å resolution. In this case, the space group was the same as observed before [13], and despite the lower resolution, some parts of the structure were indeed better resolved than in the d-P2 structure. The deprotonation of the fatty acid-coordinating Arg side chain is clearly visible in the electron density maps (Figure 1c). Figure 1d gives an example of the quality of electron density in d-P2, and the deuterium atoms can be seen even in the 2F_o_-F_c_ maps. When discussing details of hydrogen bonding and side chain conformation below, all the features are visible in the ultrahigh-resolution structures of both d-P2 and h-P2.

Anisotropy analysis of the structure (Figure 1e) shows an overall directionality of atomistic anisotropy, which could be related to functional open-close movements in the protein. Opening of the β barrel is thought to be central to ligand entry and egress, and could play a role in lipid membrane binding [18].

### 2.2. Side chain-backbone interactions

C-H…O hydrogen bonds are generally recognized to be an integral part of protein structure, especially in β sheets. Main-chain C-H…O interactions are common features of the β sheets of P2. In addition to main-chain C-H…O bonds, a number of C-H…O hydrogen bonds between side chains and the main chain are detectable. Two examples of this in P2 are provided in Figure 2.

**Figure 2.**
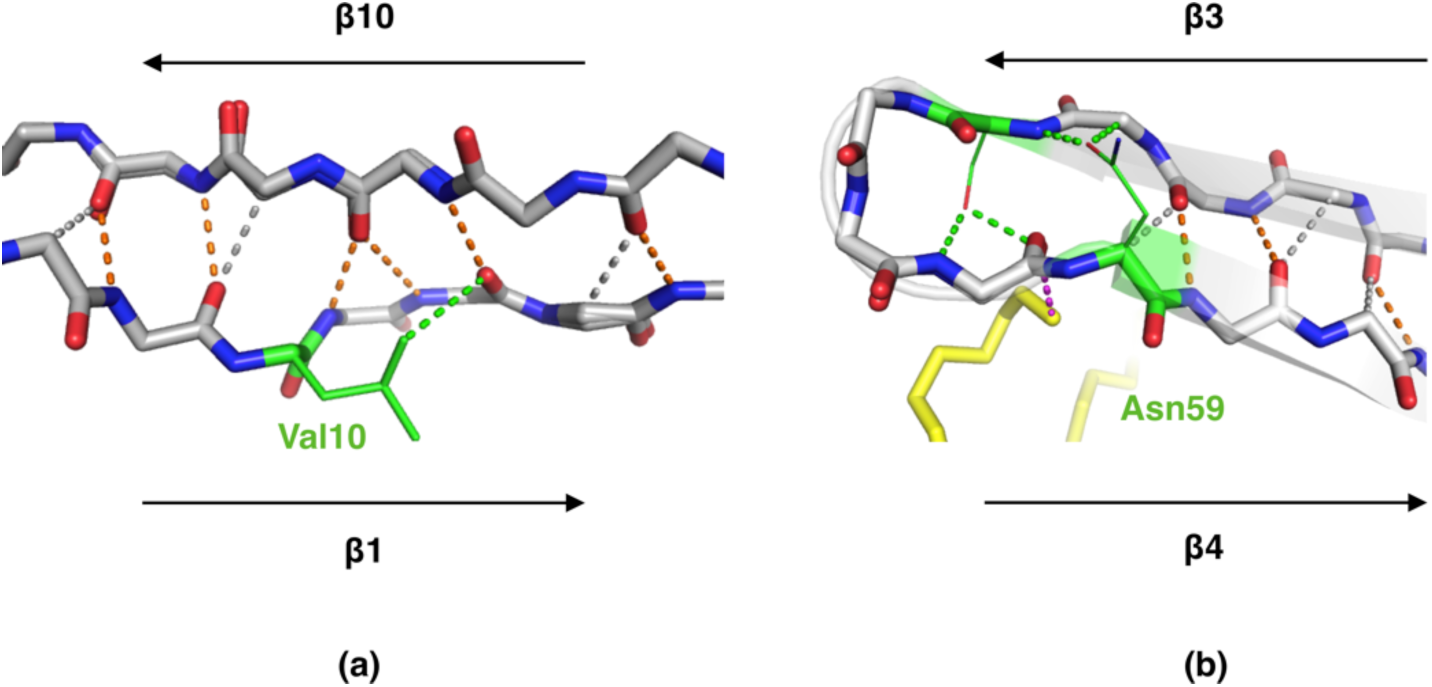
Examples of side chain involvement in β sheet hydrogen bonding. (**a**) A bump in strand β1 is observed around residue 11. A closer inspection reveals a breakdown of β sheet structure, with two NH groups H-bonding to a single carbonyl moiety and the side chain of Leu10 providing a C-H…O bond to the carbonyl group of residue 12. (**b**) Near the β3-β4 loop, which is part of the portal region and predicted to open upon ligand exchange/membrane binding, the interactions near the loop involve side chains and the fatty acid ligand (yellow), although the backbone remains in an extended β conformation. Regular hydrogen bonds are shown in orange, main-chain C-H…O bonds in gray, and bonds involving side chains in green.

C-H…O bonds between side chains and main-chain carbonyl moieties are also frequently observed at the ends of secondary structure elements (Figure 3). These interactions could be considered elements of secondary structure capping in protein structures.

**Figure 3.**
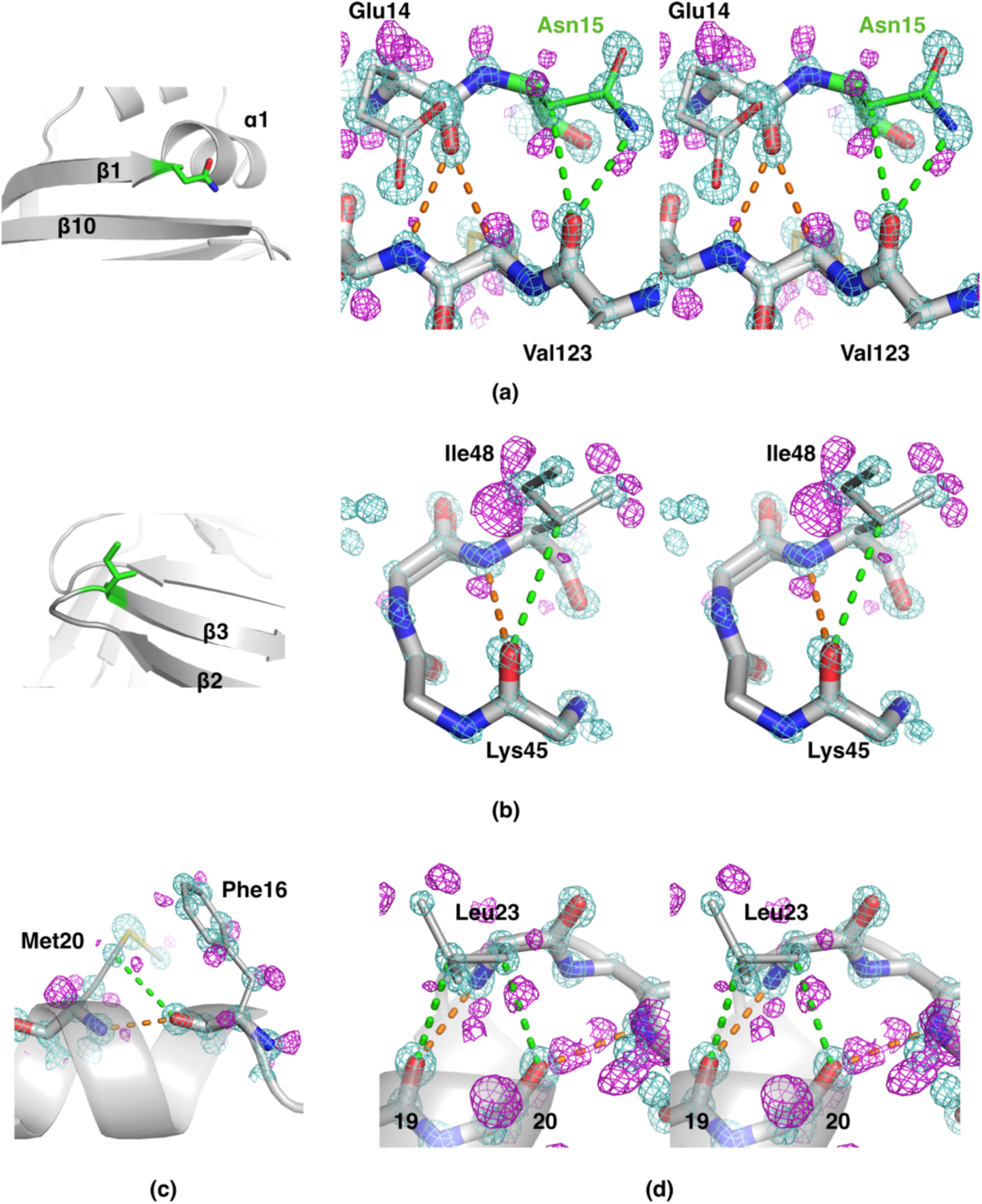
Details of protein backbone interactions from sub-atomic resolution electron density maps. Carbonyl groups in d-P2 secondary structures regularly interact with side-chain C-H groups through C-H…O hydrogen bonds. The 2F_o_-F_c_ map (cyan) in each panel is countoured at 4.0 σ, and the F_o_-F_c_ map (magenta) at 2.7, 2.5, 2.5, and 2.2 σ for panels (a)-(d), respectively. The maps were calculated after refining the structure without hydrogen atoms. Main-chain hydrogen bonds are shown in orange and hydrogen bonds involving side chains in green. (**a**) Left: Asn15 (green) is located at the C-terminal end of strand β1, just before helix α1 starts. Right: Stereo view of the interactions between the Asn15 side chain and strand β10. (**b**) The β2-β3 turn presents Cβ-H…O hydrogen bonding between Ile48 (green in the left panel) and the main-chain carbonyl of Lys45. (**c**) C-H…O bonding within helix α1, involving the side chain of Met20. (**d**) At the end of helix α1, Leu23 caps the helix through a double C-H…O interaction to the previous turn of the helix.

At the C-terminal end of strand β1, between strands 2 and 10, the strand turns away, but the β-sheet-like interactions are continued further with strand 10, through an extended conformation of the Asn15 side chain (Figure 3a). The side chain makes N-H…O and C-H…O bonds with the carbonyl oxygen of Val123 on strand β10. The sequence alignment of all human FABP family members, 12 in total, highlights that Asn15 is fully conserved, apart from human FABP5, in which it is a glycine [13]. This observation suggests that a similar “extension” of strand β1 may be a common property of FABP family members. Another location of C-H…O bonds involving side chains concerns tight β turns (Figure 3b). While the backbone carbonyl group of residue i H-bonds to the NH group of residue i+3, it also makes a C-H…O bond to the side chain of the same residue (i+3) in approximately half of the turns. In the remaining cases, the carbonyl group has a different conformation and contacts a water molecule in addition to NH(i+3).

In α helices, the carbonyl group at position (i) can be involved in a C-H…O bond to the Cβ or Cγ of a side chain in the next turn of the helix (residue i+4 or i+3) (Figure 3c). For example, the side chain of Leu23 makes two C-H…O bonds, to the backbone carbonyls of residues 19 and 20, to cap helix α1 at its C-terminal end (Figure 3d). These interactions are similar to those reported earlier in helical structures [19].

### 2.3. Unconventional side chain conformations in P2

The high resolution of the diffraction data for human P2 allows to detect unusual conformations in amino acid side chains. In fact, a few residues were listed as outliers by Molprobity, and a careful inspection confirmed the correctness of the model with respect to electron density. Notably, three arginine residues in P2 are non-planar in their guanidinium groups. In addition, Phe16 and Trp97 deviate significantly from planarity (Figure 4a).

**Figure 4.**
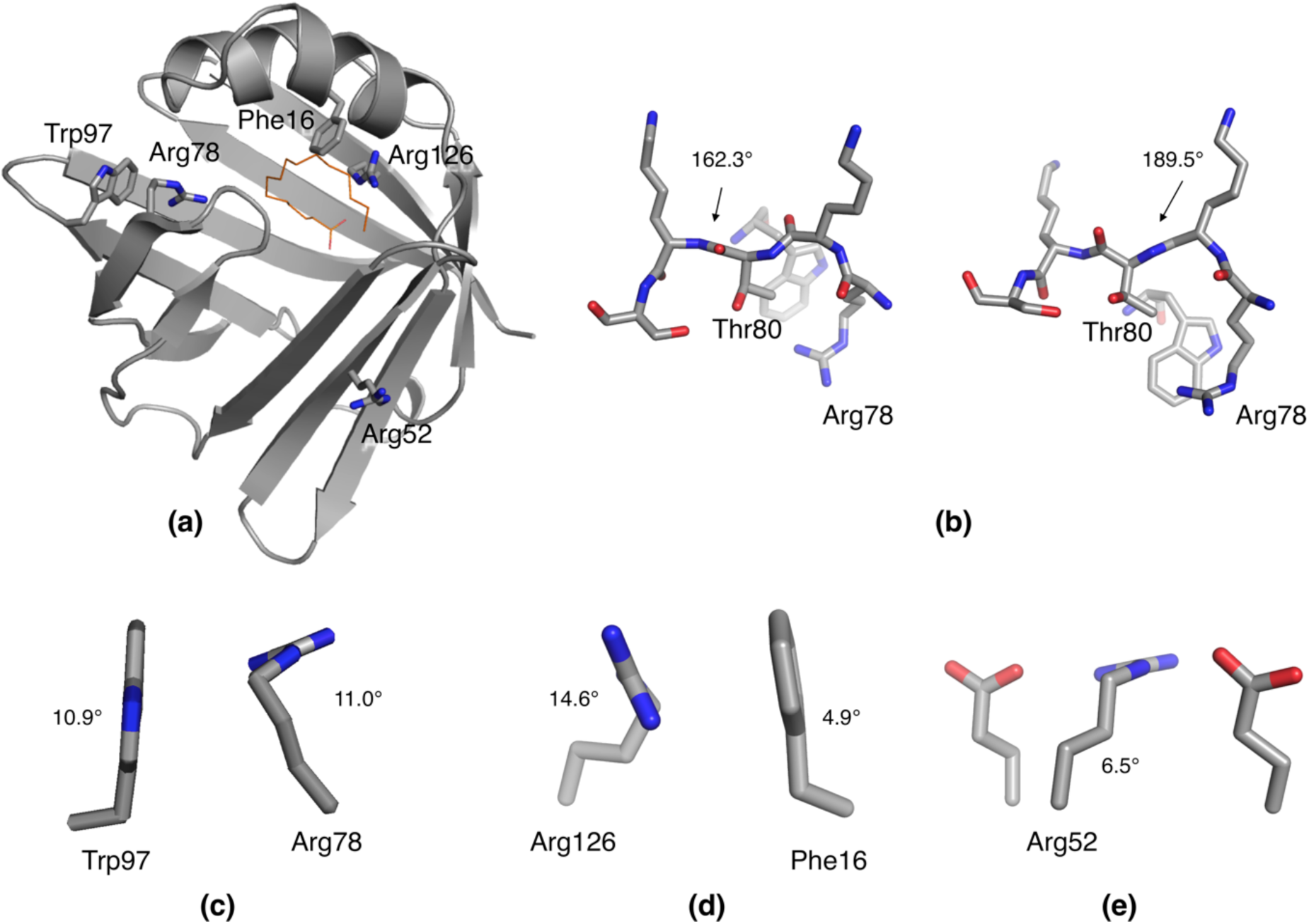
Unconventional conformations are revealed at ultrahigh resolution. (**a**) Location of the distorted side chains in the d-P2 3D structure. Note how Arg78/Trp97 and Phe16/Arg126 are in close contact. (**b**) Peptide bond distortion on both sides of Thr80 leads to its insertion deeper into the structure, interacting with Arg78 and Trp97. The ω angles for the two adjacent bent peptide bonds are shown from two slightly different views. (**c**) Packing of Trp97 against Arg78. Both side chains are strong geometric outliers due to non-planarity. (**d**) Arg126 packs against Phe16, and both residues show loss of planarity in the side chain. (**e**) Arg52 is twisted, apparently to optimize its conformation between two Glu residues.

For quantifying the analysis, the following “bending angles” were measured to reflect distortion from expected planarity. For Arg residues, the torsion angle Cδ-Nε-Cζ-Nη1 (expected 0°), for Phe, the angle [180°-(Cβ-Cγ-Cζ)] (expected 0°), for Trp, the pseudo-torsion angle Cβ-Cγ-Cε2-Cζ3 (expected 0°), and for Tyr, the sum of the angles [180°-(Cβ-Cγ-Cζ)] and [180°-(Cβ-Cζ-Oη)] (expected 0°). At high resolution, some peptide bonds deviate from planarity, the ω angle being different from the theoretical 180° (Figure 4b). These are the angles cited in the Figures.

Arg78 resides in the portal region, at the turn between strands β5 and β6, where it is engaged in a salt bridge with Asp76. In our previous study, Arg78 was one of the main residues interacting with the lipid membrane during coarse-grained MD simulations [13]. In the current structure, the side chain of Arg78 is twisted such that it can make a better salt bridge; on the other hand, a non-planar aromatic residue, Trp97, is stacked edge-to-face with Arg78 (Figure 4c). Trp97 interacts with the bent guanidinium group of Arg78 with both of its rings, possibly stabilizing this unexpected conformation. This local strain may be related to conformational changes taking place when P2 binds to a membrane surface and/or the portal region opens for fatty acid release. Such opening has thus far been observed in atomistic MD simulations and SAXS experiments [12,18].

Arg126 is one of the two arginine residues in P2 directly coordinating the carboxyl group of the bound fatty acid [13,14,18]. In h-P2, this residue was protonated, while Arg106 was deprotonated (Figure 1c). In the 0.72-Å d-P2 structure, the guanidinium group of Arg126 is a geometrical outlier, standing out at >8 σ deviation in the Molprobity analysis. A stacking interaction between Phe16 and Arg126 can be observed, and Phe16 is non-planar, with Cβ out of the phenyl ring plane. The guanidinium group of Arg126 twists so it can make face-to-face stacking with Phe16 (Figure 4d); this, on the other hand, will enable it to make optimal interactions with the ligand fatty acid. Whether this strained conformation is a unique property of the liganded state, cannot be answered at this time, since all crystal structures of human P2 thus far contain bound fatty acid.

Arg52 is on the surface of P2, making salt bridges with the side chains of Glu54 and Glu61. Its guanidinium group is well-defined an non-planar. Judging from the arrangement, the twist is linked to the formation of optimal interactions to the two acidic residues, between which Arg52 is sandwiched (Figure 4e).

Taken together, the geometric outliers observed in human P2 at high resolution can be justified based on the chemical environment. In addition, intriguing concerted bending of side chains away from planarity can be seen in cases, where the guanidium group of an Arg residue stacks against an aromatic residue. Such interactions are abundant and possibly functional in protein structures [20], and it is possible that the observations here have a more general relevance for such side chain arrangements in folded proteins.

It should be noted that these unconventional conformations will result in outliers in structure quality analysis, even though they are definitely based on real features of the high-quality experimental electron density map. Hence, a structure with outliers refined at ultrahigh resolution is partially beoynd standard structure validation algorithms, which are based on idealized geometry libraries. Too tight validation criteria may discourage crystallographers to utilize the full information available in ultrahigh-resolution crystallographic data.

### 2.4. Distorted side chain planarity in other high-resolution structures

To further assess the importance of high resolution to detect geometric outliers, selected ultrahigh-resolution protein crystal structures from the PDB were observed. In general, essentially all of such structures contain some distortions, especially bending of aromatic side chains. For example, lysozyme is a general model protein for structural biology and biochemistry, and its 0.65-Å structure [21] reveals a number of side chains with distorted planarity. The protein with the highest-resolution structure to date, crambin, also presents a twisted guanidinium group in Arg17 [5]. In cases where ultrahigh-resolution data are available from related proteins, such features appear to be conserved within protein families, suggesting they are not merely crystallographic artifacts but true properties of the proteins, likely related to functional mechanisms due to the local strain they bring about. These aspects will be discussed below.

As small proteins with a reltively large proportion of conserved aromatic residues, SH3 domains are represented in the PDB by many atomic-resolution structures. We looked at 5 SH3 domain structures, refined at <1.0 Å resolution. The structures used were PDB entries 1zuu (0.97 Å), 1tg0 (0.97 Å), 4hvw (0.98 Å), 2g6f (0.92 Å), and 2o9s (0.83 Å) [22–26]. A conserved aromatic planarity outlier is present in all structures analyzed, close to the peptide ligand-binding site; bending is most severe for a Tyr, but also the Phe residues at this position show the same phenomenon (Figure 5). This finding strongly suggests functional relevance for this structural anomaly in SH3 domains.

**Figure 5.**
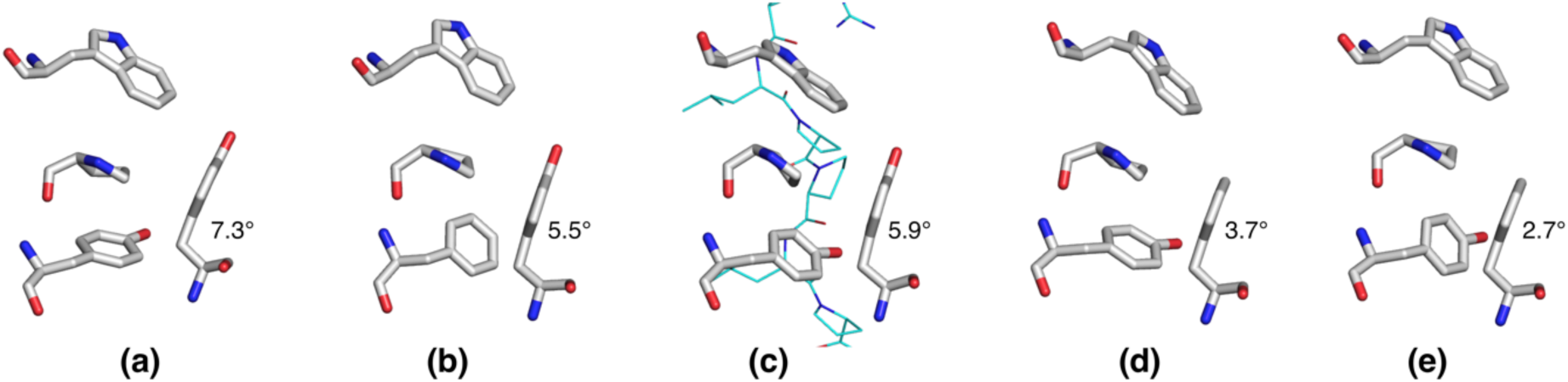
A conserved bent aromatic residue in SH3 domains. Shown is the peptide-binding site, composed of several conserved aromatic residues, of the atomic-resolution SH3 domain structures from (**a**) yeast Bzz1, (**b**) yeast Bbc1, (**c**) chicken c-Src bound to a peptide ligand, (**d**) rat betaPIX, and (**e**) human ponsin. The bending angle of the outlier aromatic residue (bottom right) is shown for all structures.

When specifically considering the function of P2, which needs to involve opening/conformational changes in the portal region, the current data provide novel ideas for the entire FABP family. The anisotropy of P2 has clear localization and direction, and these together hint at flexibility in the portal region corresponding to eventual open/close motions (Figure 1e). Opening of the portal region is a common feature proposed for the entire FABP family, and we recently observed such conformational changes in extended MD simulations [12,18]. The concentration of bent side-chain and main-chain conformations close to each other in space, especially around Arg78, is likely to be relevant for membrane or fatty acid binding-related conformational changes in P2. Since the whole FABP family is likely to share a similar conformational change related to ligand binding, it is very important to note that essentially identical deviations from ideal geometry between P2 and FABP3 [27] can be observed in their respective ultrahigh-resolution structures (Figure 6a-c).

**Figure 6.**
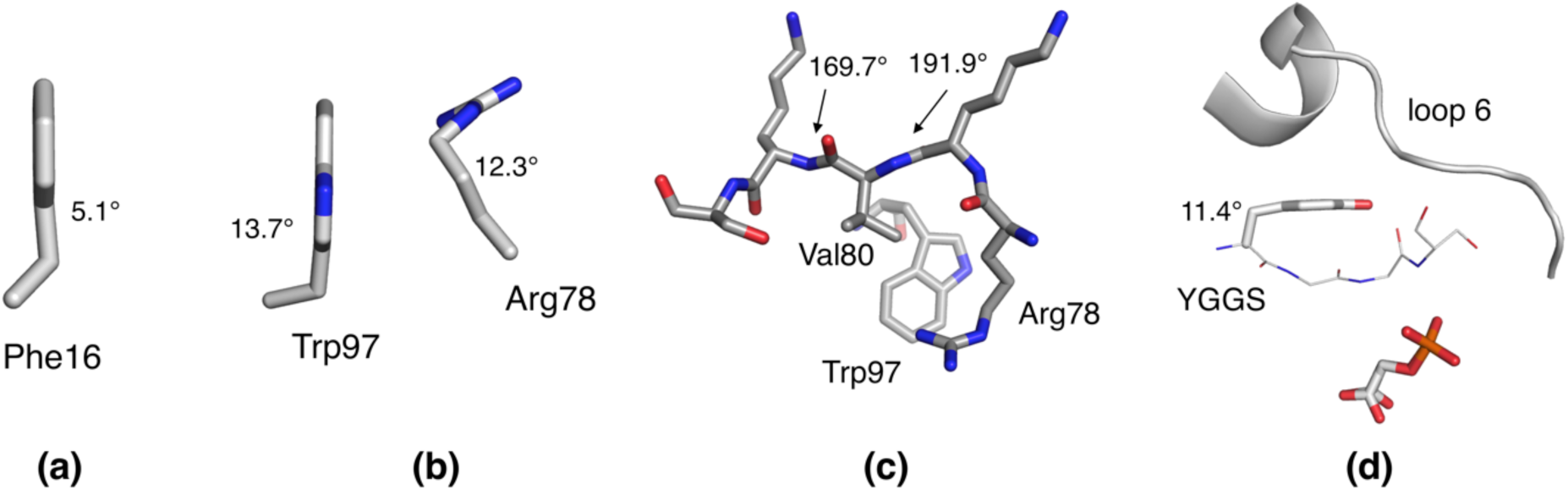
Additional geometric outliers. Compared to human P2, FABP3 shows essentially identical bending of (**a**) Phe 16, (**b**) the Trp97-Arg78 unit, and (**c**) the peptide bonds before and after residue 80. (**d**) Tyr210 is severely bent in the liganded closed structure of TIM [28].

Finally, triosephosphate isomerase (TIM) is an enzyme, for which a huge body of research data is available, including sub-atomic resolution crystal structures in complex with active-site ligands. A conserved motif in TIM, YGGS, is relevant for conformational changes and subtrate/product binding. The Tyr residue of this motif in the 0.83-Å crystal structure of TIM from *Leishmania mexicana* [28] is severely bent, and it is likely that this strain is functionally relevant (Figure 6d). Opening of the so-called loop 6, which is in close contact with Tyr210, results in the loss of the bent conformation, which can be seen at high resolution *e*.*g*. in the structure of TIM from *Plasmodium falciparum* [29].

### 2.5. Deuteration as a tool for crystallization

Proteins in biological environments are hydrogenated, apart from the rare, artificial conditions of perdeuterated protein production in culture. Isotope effects are often observed in *e*.*g*. enzymatic reactions, when comparing performance in normal and heavy water. Comparisons between deuterated and hydrogenated protein structures are important in assessing the reliability and relevance of neutron crystal structures to biological function, and they can be used to shed light on enzymatic catalysis.

Together with our earlier experiments on CNPase [30], our study suggests that deuteration could be used as a means of crystal quality optimization, even when neutron experiments are not planned. Perdeuterated CNPase produced crystals diffracting to atomic resolution and allowing to decipher protonation states and orientations of water molecules in the active site; such resolutions were never reached with the same construct without deuteration [30]. Here, human P2 crystallized in deuterated form, in D_2_O-based buffer, provided a new space group diffracting even better than the one optimized for h-P2. Several explanations can be thought of as causes of such differences; one could be shorter hydrogen bonds formed by deuterium. In addition, the recent unpublished ultrahigh-resolution structures of perdeuterated rubredoxin [6] support these findings. It might be possible to simply use D_2_O in crystallizing hydrogenated proteins, when either optimizing or screening for crystallization conditions. Perhaps deuterated conditions could be used to trap enzymatic reaction intermediates *in crystallo*. However, deuteration can can lead to changes in enzyme active-site structures *per se* [1]. Systematic studies on these aspects are clearly warranted.

## 3. Materials and Methods

### 3.1. Perdeuteration and protein purification

For production of perdeuterated human P2, the facilities at the Deuteration Laboratory, ILL in Grenoble, France were used as described [15]. Hydrogenated P2 was purified using established protocols [14].

d-P2 was analyzed by CD spectroscopy on a Chirascan Plus instrument (Applied Photophysics). CD spectra were measured at 0.25 mg/ml in a 0.5-mm cuvette. Thereafter, a thermal melting experiment was also carried out using the same instrument. For comparison, the same experiments were done using h-P2, as described earlier [13].

### 3.2. Crystallization

Perdeuterated P2 was used for growing large crystals for neutron diffraction experiments. In the course of this procedure, a large crystal representing a new crystal form was used for X-ray data collection. This crystal was grown using hanging-drop vapour diffusion equilibrating over a mother liquour consisting of 30% PEG6000 and 0.1 M sodium citrate (pD 4.25) in 90% D_2_O. Crystal growth was carried out at +8°C over a period of several months with intermittent feeding with fresh protein. The final size reached approximately 1.0×0.5×0.1 mm^3^.

Hydrogenated P2 was crystallized as previously described for the 0.93-Å structure [13]. The crystal used for data collection was grown against 100 mM sodium citrate (pH 5), 24% PEG 6000.

### 3.3. Data collection and processing

A single large large crystal of d-P2 was picked up in a loop, after a brief soak in cryoprotectant (mother liquour supplemented with 20% PEG200). The crystal was flash-cooled in liquid nitrogen. Diffraction data were collected at 100 K on the PETRAIII/DESY synchrotron radiation beamline P11 [31]. Data were processed using XDS [32]. For data collection from a h-P2 crystal with a similar procedure, beamline P13 at EMBL/DESY [33] was used. The diffraction images for both crystals can be found on the zenodo.org server [16,17].

### 3.4. Structure solution and refinement

The structure was solved by molecular replacement, using the 0.93-Å structure of P2 [13] as a search model. Refinement was carried out using phenix.refine [34] and model building using coot [35]. During refinement, hydrogen atoms were added (either as D or H), and all atoms except hydrogen were refined anisotropically. The structure was validated using Molproblity [36] and PARVATI [37]. Structure analyses and figure preparation were carried out using PyMOL [38], ccp4mg [39], and UCSF Chimera [40]. The coordinates and structure factors were deposited at the Protein Data Bank with entry codes 6S2M (d-P2) and 6S2S (h-P2).

## 4. Conclusions

The refinement of the human P2 structure at 0.72-Å resolution reveals a number of features only observable at such high resolution. These include new details on main-chain interactions, side chain conformations, and different types of hydrogen bonding. However, for accurate hydrogen/deuterium positioning of more atoms, neutron diffraction data would be required. Our earlier attempts at this produced an incomplete lower-resolution dataset [15]. On the other hand, the unexpected side chain conformations discussed above did not catch our attention during the refinement of the previous 0.93-Å structure of the hydrogenated P2 protein [13]; in retrospect, most of them can be identified in the earlier structure. Hence, for accurate descriptions of non-ideal conformations and local strain in crystal structures, it is clear that resolutions well below 1.0 Å are required. One must also have a keen eye on looking for such anomalies, instead of trying to fit every residue into a geometric ideal norm. Such data, currently only attainable through X-ray crystallography, will give novel insights into the detailed mechanisms of protein function in biological systems, even conserved across entire protein families.

## Author Contributions

Conceptualization, S.L. and P.K.; Methodology, S.L. and P.K.; Investigation, S.L. and P.K.; Data Curation, P.K.; Writing – Original Draft Preparation, P.K.; Writing – Review & Editing, S.L. and P.K.; Visualization, P.K.; Supervision, P.K.; Project Administration, P.K.; Funding Acquisition, P.K.

## Acknowledgments

We wish to thank Ravi Yadav for crystallizing h-P2. We are grateful for beamtime and excellent synchrotron beamline support at PETRAIII/DESY/EMBL crystallography beamlines.

## Sample Availability

Samples of the P2 protein and its expression vector are available from the authors upon reasonable request. Raw diffraction images are available through zenodo.org, and the refind coordinates and structure factors from the Protein Data Bank.

